# Direct measurement of the aerotactic response in a bacterial suspension

**DOI:** 10.1101/2022.03.09.483614

**Authors:** J. Bouvard, C. Douarche, P. Mergaert, H. Auradou, F. Moisy

**Affiliations:** Université Paris-Saclay, CNRS, FAST, 91405, Orsay, France; Université Paris-Saclay, CEA, CNRS, Institute for Integrative Biology of the Cell (I2BC), 91198, Gif-sur-Yvette, France

## Abstract

Aerotaxis is the ability of motile cells to navigate towards oxygen. A key question is the dependence of the aerotactic velocity with the local oxygen concentration *c*. Here we combine simultaneous bacteria tracking and local oxygen concentration measurements using Ruthenium encapsulated in micelles to characterise the aerotactic response of *Burkholderia contaminans*, a motile bacterium ubiquitous in the environment and present in living organisms. In our experiments, an oxygen gradient is produced by the bacterial respiration in a sealed glass capillary permeable to oxygen at one end, producing a bacterial band travelling towards the oxygen source. We compute the aerotactic response *χ*(*c*) both at the population scale, from the drift velocity in the bacterial band, and at the bacterial scale, from the angular modulation of the run times. Both methods are consistent with a power-law *χ ∝ c*^*−*2^, in good agreement with existing models based on the biochemistry of bacterial membrane receptors.

## I. INTRODUCTION

Motile bacteria have the ability to bias their swimming in response to a gradient of chemicals in order to move towards an optimum environment [1]. Aerotaxis [2, 3], the specific response to oxygen, is a common feature of a large number of bacterial species [4–12]. This phenomena was first highlighted by Engelmann [2] in 1881 when he used bacteria to spot sites of oxygen produced by photosynthetic algae.

Aerotaxis plays a key role in many biological processes such as biofilm formation [13–16], or in the bacterial distribution in the rhizosphere, the soil environment that is under the direct influence of plant roots, where the respiration and the metabolic activities of the roots create oxygen gradients [17]. In the field of health, aerotaxis was recently suspected to influence the colonisation of epithelial surfaces [18]. This ability to respond positively or negatively to oxygen gradients also opened the possibility to use bacteria to target hypoxic tumours that are resistant to drugs [19, 20]. It was observed that not all bacteria swim toward a high level of oxygen like the facultative anaerobe *Escherichia coli* [5] or other strictly aerobic bacteria [4, 6]. Indeed, some strains avoid microenvironments too rich in oxygen [7], while others are microaerophilic and prefer a low but finite level of oxygen [8–12]. A good qualitative description of aerotaxis is also important to prevent the quality deterioration of food [21], or to quantify bacterial flux towards regions contaminated by heavy metals [8, 13, 16].

Even if the aerotactic behaviour of bacteria has been widely studied, questions concerning the exact dependence of the aerotactic response with the environmental conditions are still open. The cellular basis of aerotaxis in swimming bacteria is the modulation of their flagellar activity in response to changes in the oxygen level sensed by their receptors [22–26]. This translates into a net aerotactic velocity ***v***_***a***_ which writes, in the limit of small oxygen gradient **∇***c*,

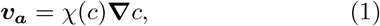

where *χ*(*c*) is the aerotactic response. As a result, aerotactic bacteria that swim towards oxygen-rich regions are able to collectively migrate, forming a travelling band at the population scale.

Travelling bands have been widely investigated for various chemoattractants [3, 27–29] - we restrict here the discussion to aerotaxis. Their dynamics is described by the Keller-Segel equations for the bacterial concentration *b*(**x**, *t*) and oxygen concentration *c*(**x**, *t*), that write in 1D [30]

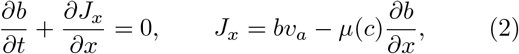

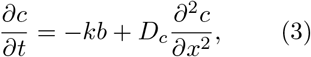

with *J*_*x*_ the bacterial flux. The first contribution in *J*_*x*_ is the aerotactic flux, with *v*_*a*_ = *χ*(*c*)*∂c/∂x* the 1D form of the aerotactic velocity (1), and the second contribution is the diffusive flux, with *μ*(*c*) the diffusion coefficient due to random swimming (also called random motility coefficient [31–33]). The second equation describes the dynamics of the oxygen concentration, with *k* the oxygen consumption rate by bacteria and *D*_*c*_ the oxygen diffusivity.

The dynamics of the bacterial migration critically depends on the dependence of *χ* with *c*. A minimum requirement for the formation of such a travelling band is a decrease of *χ*(*c*) as *c*^*−*1^ or steeper [30]. Models based on the biochemistry of bacterial membrane receptors predict an aerotactic response in the form [31, 34]

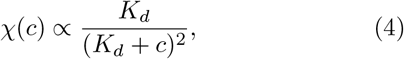

with *K*_*d*_ the dissociation constant of the considered chemoattractant from its receptor, i.e., the concentration of the chemoattractant such that half of the receptors are bound. Early experiments by Brown and Berg on the chemotaxis of *E. coli* [35] are consistent with this form. For small molecules such as dioxygen, *K*_*d*_ is usually much smaller than the ambient concentration [36], yielding an effective scaling *χ* ∝ *c*^*−*2^.

It is possible in principle to infer the aerotactic response *χ*(*c*) from the dynamics of the bacterial concentration profile evolving in a 1D oxygen gradient, provided that the local oxygen concentration *c*(*x*) and *μ*(*c*) are known. Recently, microfluidic setups where the oxygen gradient is fixed by imposing the concentration at the boundaries were developed [4, 37–40]. In this configuration, if the variations of *μ* with the oxygen concentration can be neglected, the steady solution of Eq. (3) is an exponential profile *b*(*x*) resulting from the balance between the diffusive and the aerotactic fluxes, from which *χ* can be obtained. However, accurate measurements can only be obtained for decay lengths falling between a few micrometers up to the channel size of a fraction of millimetres, limiting in turn the range of parameters that can be tested. These studies tend to show the existence of a “log-sensing” regime, *χ*(*c*) ∝ *c*^*−*1^, for moderate oxygen gradients (the corresponding aerotactic velocity is *∂*(ln *c*)*/∂x*), although other scalings like Eq. (4) cannot be excluded over wider range of oxygen gradients.

The recent development of O_2_ bio-compatible sensors offers now the possibility to perform simultaneous measurements of oxygen concentration and bacterial velocity statistics. In this study, we use Ru-micelles, which consist of Ruthenium (an oxygen-quenched fluorophore) encapsulated in micelles [5, 41], to measure the oxygen concentration in a bacterial suspension enclosed in an elongated glass capillary. The glass capillary is closed at one end but permeable to air on the other, allowing an unidirectional oxygen gradient. The bacterial respiration lowers the oxygen concentration in the suspension, which in turn strengthens the gradient, resulting in a travelling band and a strong accumulation of bacteria near the oxygen source at large time. The simultaneous measurements of oxygen and bacterial tracks allow us to compute directly the aerotactic response *χ*(*c*), both at the population scale and at the bacterial scale. Our results suggest a robust *χ*(*c*) ∝ *c*^*−*2^ scaling, consistent with Eq. (4) for a dissociation constant *K*_*d*_ much smaller than the ambient oxygen concentration level.

## II. MATERIAL AND METHODS

### A. Experimental setup

The experimental setup is shown in Fig. 1(a). A homogeneous suspension of *Burkholderia contaminans* bacteria is enclosed in a glass capillary (Vitrotubes) of rectangular cross section (0.4 mm height, 8 mm width) and 50 mm long. The capillary is closed at one end by PDMS (polydimethylsiloxane) plug, a biocompatible cured polymer permeable to oxygen, and sealed at the other end by grease. The PDMS plug is made by dipping the capillary in liquid PDMS, resulting in a plug of (13.0 ± 0.5) mm long.

**Figure 1.**
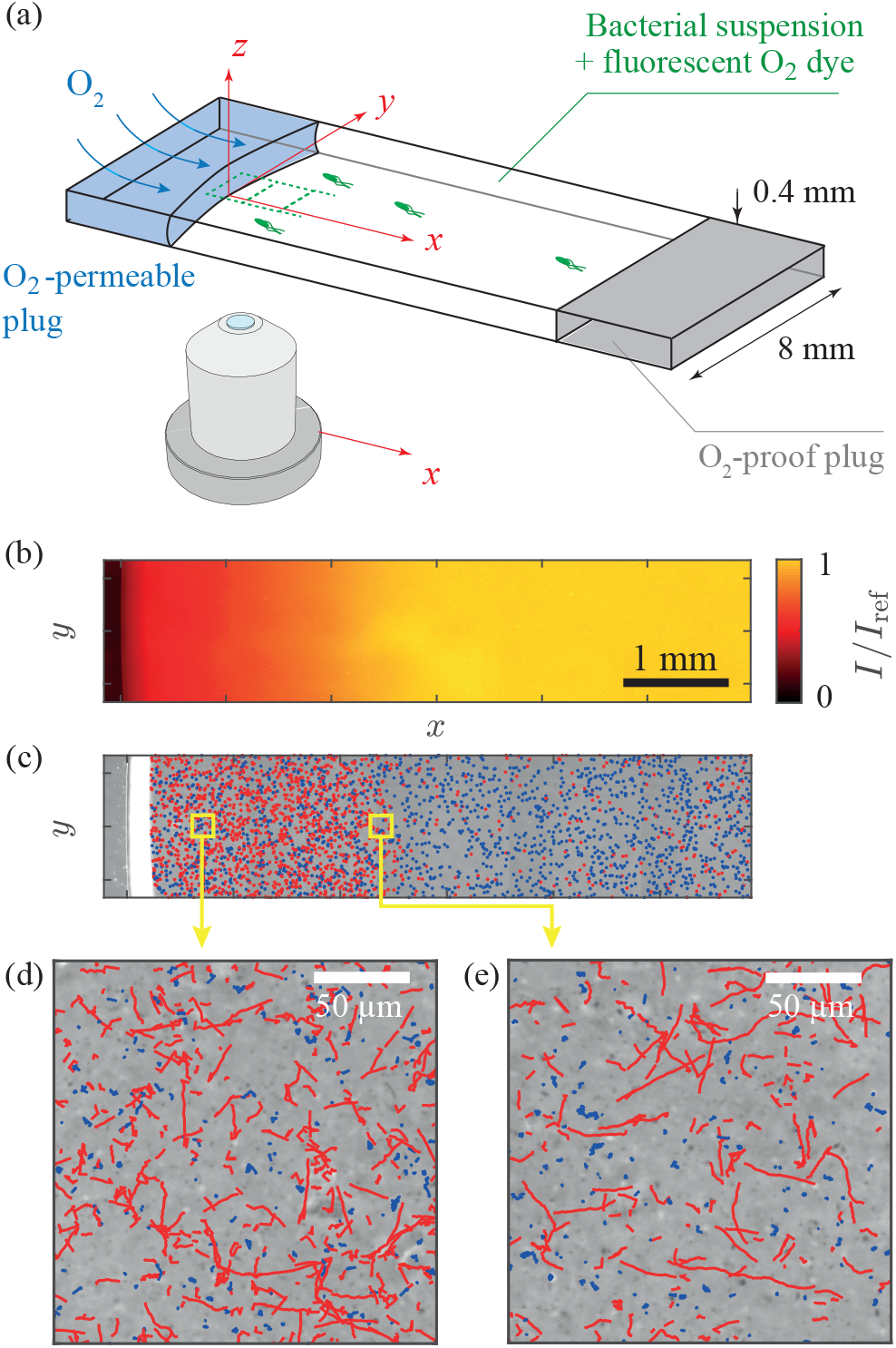
Aerotactic migration band in a glass capillary permeable to air on one side. (a) Experimental setup, made of a glass capillary of rectangular section, filled with both the bacterial suspension and the fluorescent oxygen dye. One side of the capillary is sealed with an oxygen-porous PDMS plug, while the other side is sealed with an oxygen-proof grease plug. (b) Fluorescence light intensity mapping the oxygen concentration field at large time, during the migration of the travelling band (dark: high O_2_ concentration). (c) Images of bacteria at the same time, superimposed to the bacteria tracks (non-motile and motile bacteria are respectively in blue and red). (d), (e) Magnifications at two locations, showing the nearly random tracks at large oxygen concentration and the strongly biased tracks at low oxygen concentration.

The bacterial strain of *B. contaminans* used in this study is an environmental strain isolated in the FAST laboratory. This strain was selected based on its aerotactic ability in a preliminary screening of several available strains, and characterised from the sequencing of 16S and recA gene fragments. Bacteria are grown overnight in YEB medium (Yeast Extract Beef: 5 g L^*−*1^ beef extract, 1 g L^*−*1^ yeast extract, 5 g L^*−*1^ peptone, 5 g L^*−*1^ sucrose, 0.5 g L^*−*1^ MgCl_2_) at 28 ^*°*^C in an orbital shaker running at 200 rpm up to an optical density OD_600_ = 5.5 ± 0.5. The suspensions are then diluted in YEB to the desired concentration, and 1 μmol L^*−*1^ of Ru-micelles [41] is added to the suspension. The solution is finally homogenized with an orbital vortex mixer before injection in the capillary. Experiments were repeated for three bacterial concentrations, denoted hereafter by their optical densities OD = 0.05, 0.1 and 0.2 (1 OD ∼ 1.8 ×10^6^ bact μL^*−*1^).

The capillary is located on the stage of an inverted microscope (Leica DMI4000B) equipped with a long working distance 10X objective and a Hamamatsu Orca Flash 4 camera allowing to record fields of view of (1.33 × 1.33) mm^2^ with 30 μm depth. Two imaging methods are used sequentially: fluorescence imaging, from which the oxygen concentration is computed (see Fig. 1(b) and Appendix A), and phase contrast for bacteria tracking [Fig. 1(c)]. Measurements are performed at 5 locations along the *x* axis with 10 % overlap, covering a distance of nearly 6 mm in the *x*-direction from the source at *x* = 0. The acquisition sequence is as follows: a fluorescence picture is first taken at each of the 5 locations, then a movie of 100 images at 20 Hz is acquired in phase contrast at the same locations. The sequence is repeated up to 20 times during the bacterial migration, with a time step between each scan of 3 to 10 minutes depending on the migration timescale. The maximum time lag between an oxygen measurement at a given location *x* and the bacteria tracks at the same location is less than 90 s, a value comfortably smaller than the time scale of the band migration.

### B. Bacteria tracking

From the sequences of phase contrast images, the bacterial tracks are computed using Trackmate [42] and post-processed using an in-house Matlab code. For each track *i*, we measure the two-dimensional positions **X**_*i*_(*t*) = (*X*_*i*_(*t*), *Y*_*i*_(*t*)), from which we compute the two-dimensional velocities **V**_*i*_(*t*) = *δ***X**_*i*_*/δt* using a sampling time *δt*. This sampling time is chosen such as to separate the motile and non-motile populations that are naturally present in the suspension in approximately the same proportions [red and blue tracks in Fig. 1(d) and (e)]. During the time *δt*, motile cells travel a distance ≃ *V*_0_ *δt*, with *V*_0_ ≃ 20 μm s^*−*1^ the typical swimming velocity (see Fig. 2). During the same time *δt*, non-motile cells move randomly over a distance 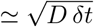, with *D ≃* 0.1 μm^2^ s^*−*1^ the thermal diffusion coefficient (as measured independently from the mean-square displacement [43]). A clear separation between the two populations is obtained by using a time interval 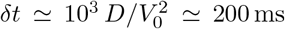. With this value, non-motile bacteria have an effective velocity 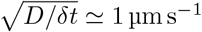 much smaller than that of motile bacteria, and can be filtered out by applying a velocity threshold at 6 μm s^*−*1^. Only motile bacteria are considered in the following.

**Figure 2.**
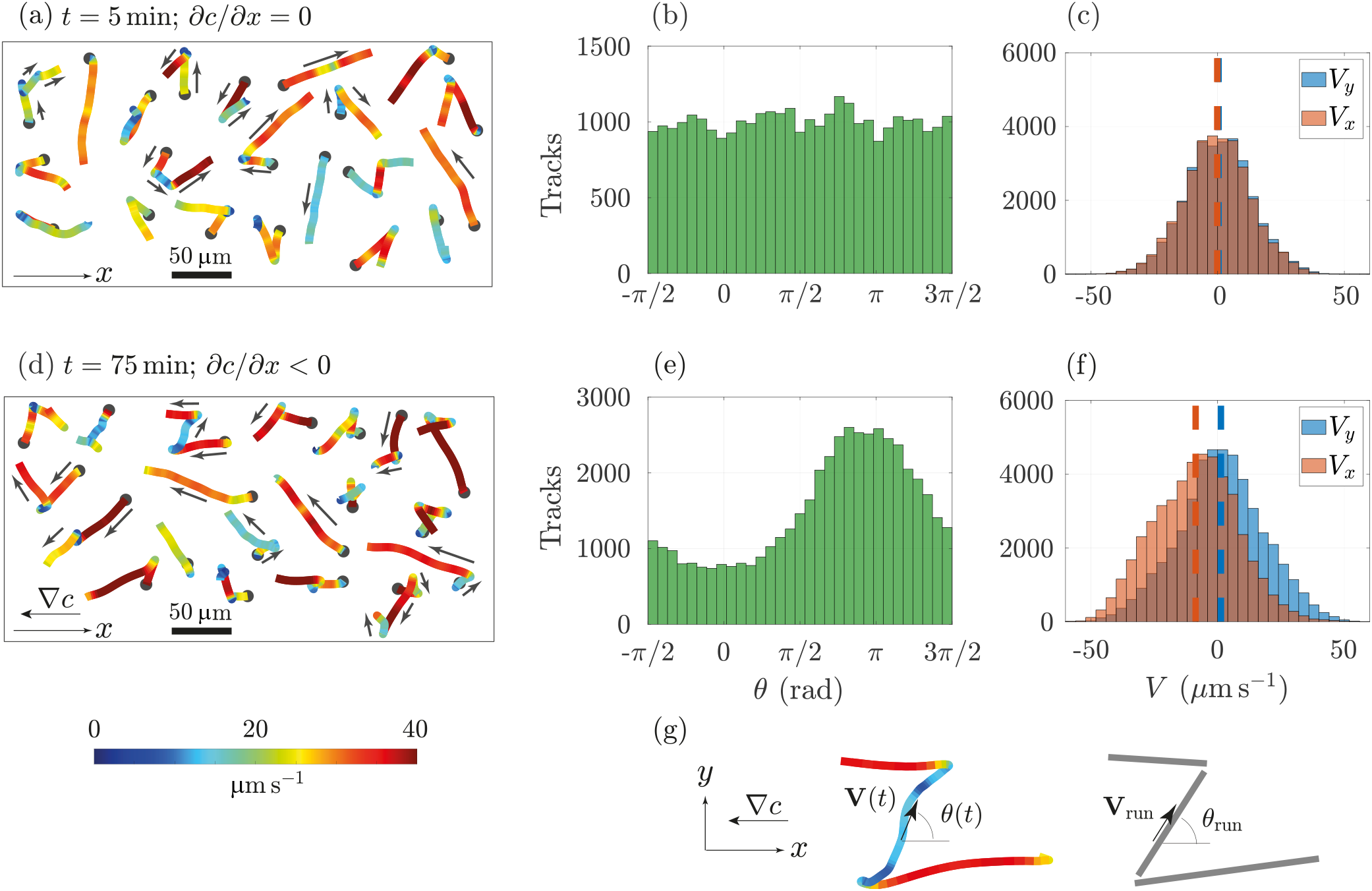
Characteristics of bacterial tracks without and with aerotaxis. Data from the experiment at OD = 0.05, at location *x* = (4500 *±* 500) μm from the oxygen source. (a), (b), (c) *t* = 5 min, while the oxygen concentration is homogeneous in the glass capillary; (d), (e), (f) *t* = 75 min, when a steep oxygen gradient is formed. (a), (d) Selection of bacterial tracks, coloured according to their instantaneous velocity. The starting point of a track is represented with a grey circle. The track durations are between 3 and 5 s. (b), (e) Histograms of the instantaneous trajectory angle *θ*. (c), (f) Histograms of the instantaneous velocity components *V*_*x*_, *V*_*y*_. (g) Segmentation of tracks into a set of runs, defining the run velocity **V**_run_ and run angle *θ*_run_ with respect to *x*.

A selection of tracks is shown in Fig. 2, without (a) and with (d) aerotaxis. These tracks display a succession of approximately straight runs, typically 10 to 100 μm, separated by rapid changes in orientation. A specificity of the swimming pattern of *B. contaminans* is the frequent occurrence of very sharp turning angles close to 180° (see Appendix B), contrarily to the smaller mean turning angle of ≃ 70° of the classical “run-and-tumble” pattern in the model bacterium *E. coli* [44, 45]. Such a “run-reverse” pattern is also encountered in some marine bacteria [46–48]. In the presence of the oxygen gradient, runs in the direction of the oxygen source (*x <* 0) are longer and more frequent. This bias is difficult to see directly in Fig. 2(a,d), but appears clearly in the distributions of velocity components and angle in Fig. 2(b,c,e,f). The instantaneous swimming angle *θ*(*t*) is defined in Fig. 2(g), with *θ* = *π* toward the oxygen source (*x <* 0). In the presence of the oxygen gradient, the distribution of angles shows a strong asymmetry, with tracks in the oxygen direction approximately 3 times more frequent than in the opposite direction. Interestingly, the distribution of *V*_*x*_ is peaked to a negative value, but its width is comparable to that of *V*_*y*_. This feature is robust at all times, with a ratio of standard deviations *σ*(*V*_*x*_)*/σ*(*V*_*y*_) ≃ 1.00 ± 0.01, indicating that the bacterial motion can be approximately modelled as the sum of an isotropic diffusion and a small deterministic drift towards *x <* 0.

We determined the diffusivity of the bacteria without aerotaxis in two setups: in a chamber fully permeable to oxygen (with a PDMS cover) and in a fully sealed capillary (capillary glass sealed with a grease plug at both ends). In the first setup, the oxygen concentration remains at saturation level at all time, while in the second it linearly decreases in time because of bacterial respiration. From the bacterial tracks, we compute the diffusivity as 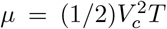, where *V*_*c*_ and *T* are determined from an exponential fit of the two-dimensional velocity correlation function (see Appendix C). We obtain *μ* (450 ± 100) μm^2^ s^*−*1^, with no significant evolution in time in both setups as long as oxygen is available. In the sealed capillary, the motility rapidly drops when oxygen is exhausted and transitions to a brownian diffusive motion. Similarly, the mean 2D velocity stays constant throughout both experiments, and only drops when the oxygen concentration reaches zero in the sealed capillary experiment.

## III. DYNAMICS OF THE AEROTACTIC FRONT

To quantify the dynamics of the aerotactic migration of bacteria towards the oxygen source, we compute the velocity statistics conditioned on the *x* position along the capillary. For this, we first define for each track *i* its average position 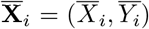 and average velocity 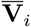. Then, we average the velocity norm and *x*-component conditioned on the averaged *x*-position,

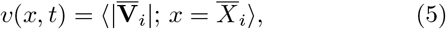

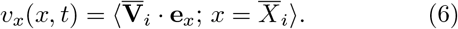

The brackets ⟨·⟩ denote the average over all tracks *i* satisfying this spatial conditioning. In practice, the conditioning is performed on a window size of width Δ*x* = 200 μm. This window size is chosen larger than the typical run length, but smaller than the characteristic length scale of the oxygen and bacterial concentration profiles. Note that the velocities in Eqs. (5)-(6) are the two-dimensional projections of the true bacteria motion. This implies that for bacteria swimming isotropically at constant (three-dimensional) velocity *V*_0_, we have *v* = *πV*_0_*/*4 ≃ 0.78*V*_0_, whereas for bacteria swimming in the microscope plane we simply have *v* = *V*_0_.

The dynamics of the migration band is depicted in Fig. 3 along with the oxygen concentration profile, the mean velocity norm *v*(*x*) and the drift velocity along the gradient *v*_*x*_(*x*). Five minutes after the sealing of the capillary [Fig. 3(a)], the oxygen level has slightly decreased but remains essentially homogeneous, and no measurable drift velocity nor variations in the bacterial concentration are detected. The migration band starts at *t* ≃ 75 min [Fig. 3(b)], when the oxygen level reaches zero at the end of the measurement area (*x* > 5 mm). In the oxygen-depleted region, a bump of bacterial concentration appears together with a strong drift velocity, *v*_*x*_ ≃ − 8 μm s^*−*1^ (negative velocity is towards the oxygen source), which is up to 30 % of the 2D mean swimming velocity. Ten minutes later [Fig. 3(c)], the bacterial band is more pronounced as it approaches the oxygen source, but it travels at a smaller velocity, *v*_*x*_ ≃ − 3.5 μm s^*−*1^. After 100 min, an asymptotic state is reached, with a strong accumulation of bacteria near the O_2_ source and *v*_*x*_ ≃ 0 μm s^*−*1^ [Fig. 3(d)].

**Figure 3.**
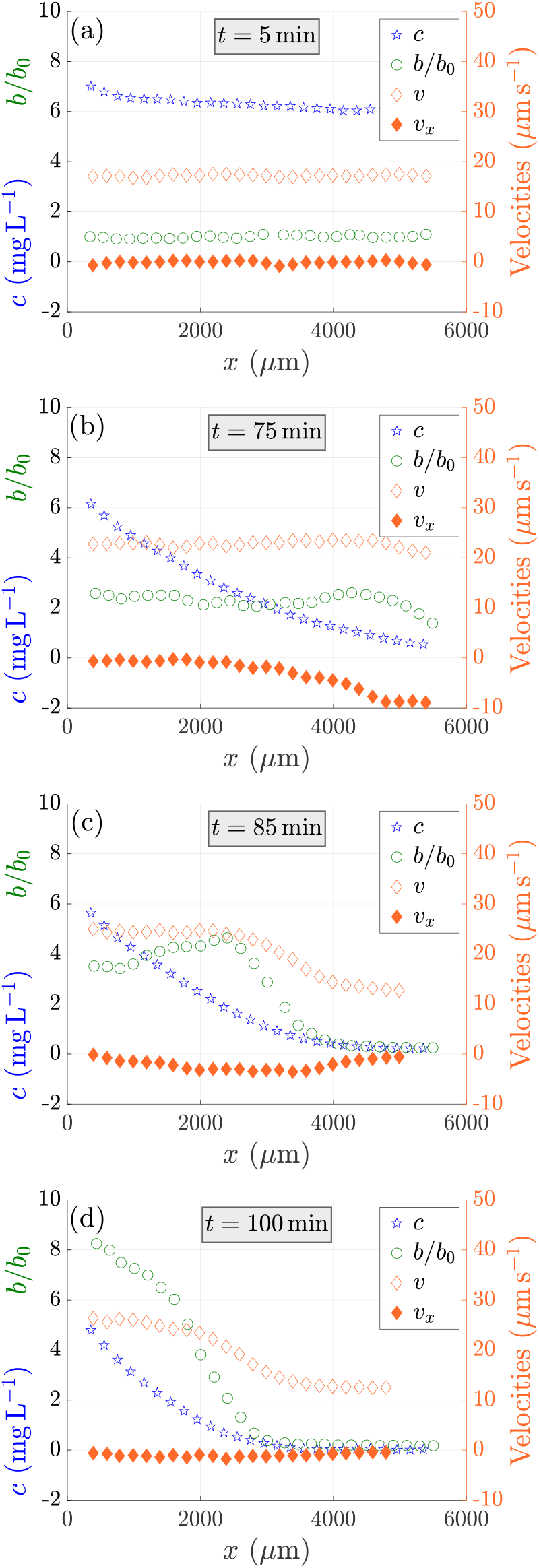
Variations of the local density of bacteria *b*(*x*) (green circles), oxygen concentration *c*(*x*) (blue stars), magnitude of the swimming velocity of bacteria *v*(*x*) (empty orange diamonds) and drift velocity along the gradient *v*_*x*_(*x*) (filled orange diamonds) as functions of the distance from the oxygen source at four different times during the migration of the bacterial band. The capillary is initially filled with an homogeneous suspension of bacteria at a concentration OD = 0.05. Time is counted from the sealing of the capillary.

The experiment was repeated with larger initial bacterial concentrations (see Appendix E). The dynamics of the traveling band is similar, but on a time scale that varies approximately as the inverse of the bacterial concentration. An example is given in Fig. 4 at OD = 0.2, clearly showing the shift of the maximum drift velocity as the migration band proceeds to the oxygen source.

**Figure 4.**
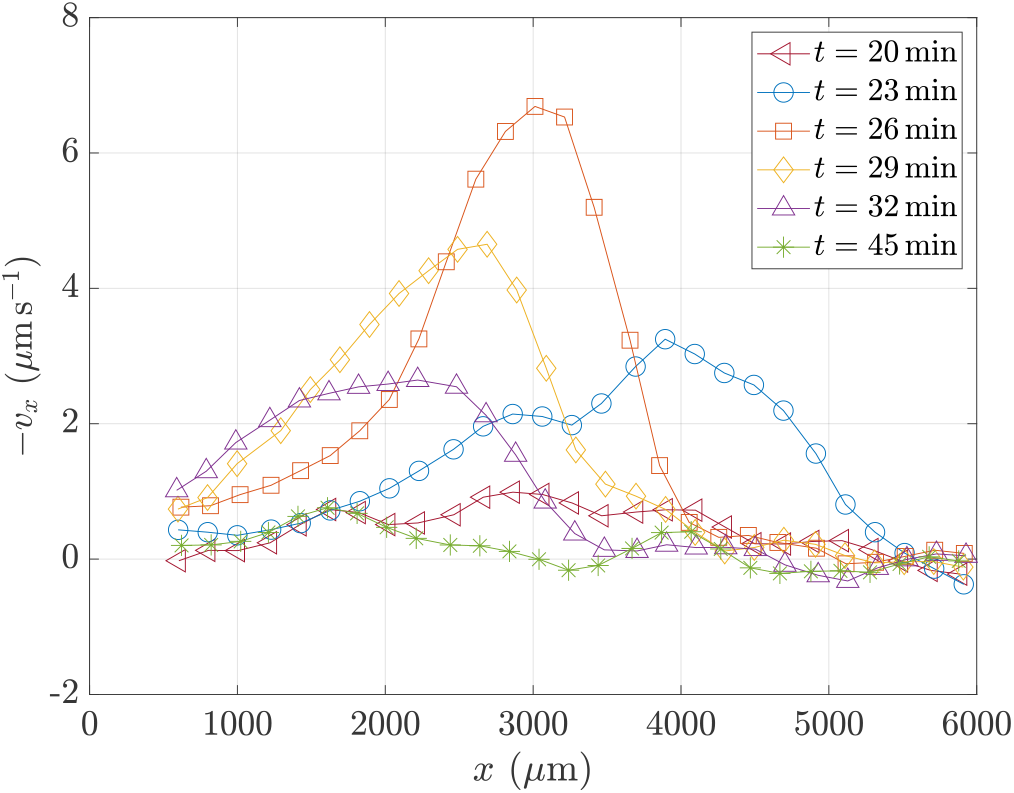
Drift velocity *−v*_*x*_(*x*) as a function of the distance from the oxygen source at different times during the migration of the bacterial band. The initial bacterial concentration is OD = 0.2.

During the migration process, a significant increase of the mean 2D velocity *v*, from 18 to 26 μm s^*−*1^, is found (Fig. 3). This increase may originate from an intrinsic dependence of *v* with the oxygen concentration level (chemokinesis), or may be an indirect consequence of aerotaxis. To discriminate between these two hypotheses, we measured the mean swimming velocity in homogeneous experiments, in the same setups as for the diffusivity measurements: in the chamber fully permeable to oxygen and in the fully sealed capillary. In both cases we found a constant mean swimming velocity, with no dependence neither on time nor on oxygen concentration. This shows that the temporal increase in mean velocity in Fig. 3 is not a response to the local oxygen concentration, but rather a consequence of aerotaxis. This increase may be due in part to a projection effect: for bacteria swimming isotropically at velocity *V*_0_, the projected mean velocity in the (*x, y*) plane is *v* ≃ 0.78*V*_0_, whereas for a strongly biased swimming in the *x*-direction the projected velocity *v* is equal to the true velocity *V*_0_. However, this projection effect alone cannot explain the observed 40% increase in velocity, suggesting that a sorting effect may also be at play: slow and fast bacteria are initially randomly distributed in the suspension, but faster bacteria first reach the oxygen source while slow bacteria remain behind in the oxygen depleted region, thereby introducing a bias in the mean velocity.

We now turn to the determination of the aerotactic coefficient *χ*(*c*) = *v*_*a*_*/*(*∂c/∂x*), where *v*_*a*_ is the aerotactic velocity. Measurements are averaged over a duration of 5 s, which is much smaller than the timescale of the evolution of the oxygen and bacterial concentration profiles. A difficulty in inferring *v*_*a*_ from the measured drift velocity *v*_*x*_ is that *v*_*x*_ is related to the *total* bacterial flux *J*_*x*_ = *bv*_*x*_, and therefore includes both the intrinsic aerotactic contribution *bv*_*a*_ and the diffusive contribution *μ*(*c*)*∂b/∂x* [Eq. (3)]. Correcting for this diffusive contribution is delicate, because the dependence of *μ* with *c*, which stems from the dependence of *χ* with *c*, is difficult to infer from the data. We therefore focus on regions where *∂b/∂x* ≃ 0, near the maximum of the bacterial density in the migrating band or close to the O_2_ source. In these regions, the diffusive flux is small and we have *v*_*x*_ *≃ v*_*a*_, so the aerotactic response is simply given by

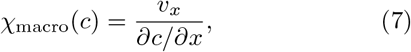

which can be directly computed from our velocity and oxygen concentration measurements. We note *χ*_macro_ this first determination of the aerotactic coefficient, as it is obtained from the drift velocity at the macroscopic level.

Figure 5 shows the aerotactic response *χ*_macro_ as a function of *c* at different times during the migration. The data is well described by a power law, *χ*_macro_ ∝ *c*^*−*2^, over the whole range of measurable oxygen concentration, from *c* = 0.6 to 4 mg L^*−*1^. The upper bound of this range is limited by the smallest measurable drift velocity, while the lower bound is limited by the resolution of the fluorescence signal. The good collapse of the data at different times on a single curve indicates that the aerotaxis response is governed by the local oxygen concentration alone, with no measurable delay, so it can be considered as approximately instantaneous in our experiment. This scaling *χ*_macro_ **∝** *c*^*−*2^ is consistent with Eq. (4) in the limit *K*_*d*_ *≪ c*. Although the dissociation constant *K*_*d*_ of oxygen is not known for *B. contaminans*, the absence of visible cutoff in *χ*_macro_(*c*) at small *c* suggests that *K*_*d*_ is much smaller than the minimal oxygen concentration measured in Fig. 5 (≃0.6 mg L^*−*1^). This is consistent with the value of *K*_*d*_ ≃0.02 mg L^*−*1^ determined for other strains like *Escherichia coli* and *Salmonella typhimurium* [36].

**Figure 5.**
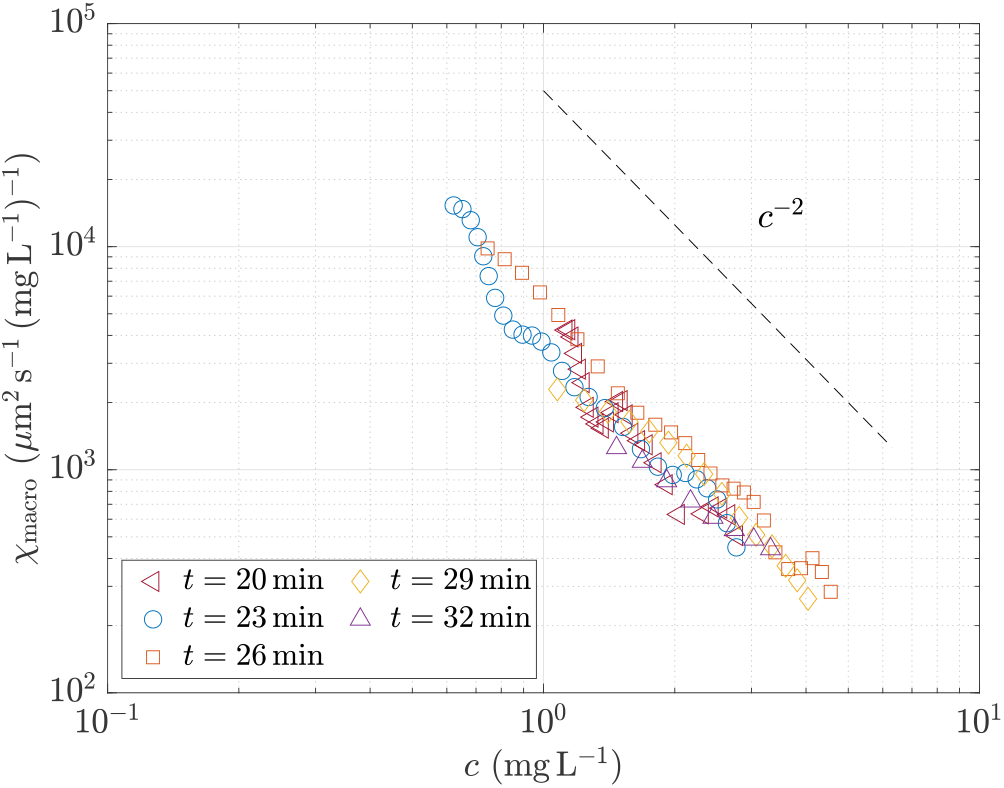
Aerotactic response *χ*_macro_ ≃ *v*_*x*_*/*(*∂c/∂x*) as a function of the local oxygen concentration *c*. The values are computed from the drift velocity *v*_*x*_(*x*) and oxygen gradient *∂c/∂x* measured in regions where *∂b/∂x* ≃ 0, for an initial bacterial concentration OD = 0.2.

We have repeated the measurement of *χ*(*c*) for the three bacterial concentrations. The curves are shown in Fig. 6, with error bars reflecting the dispersion for each OD. Here again, the data is well described by the power law *χ ≃ δc*^*−*2^, but with a coefficient *δ* that decreases as the bacterial concentration increases (see inset). This suggests that the scaling of the aerotactic response with oxygen concentration is robust, but the level of this response also depends on other parameters related to the bacterial concentration. The delay between the measurements of the oxygen concentration and the bacterial tracks cannot explain this dependence. Because of the sequential acquisition procedure (see Sec. II B), the oxygen concentration is measured approximately 1 min before the bacterial velocity. During this time lag, *c* decreases while |*∂c/∂x* | increases at a given *x*, at a rate which increases with the OD; this should increase the apparent *δ*, which does not match our observations. A second possible bias is a selection of the fastest swimming bacteria as the OD is increased; however, this sorting effect would also yield an increase of the apparent, which again is not compatible with our data. We can conclude that the observed decrease of *δ* with the OD is a genuine feature of the aerotactic response of this strain, that may

**Figure 6.**
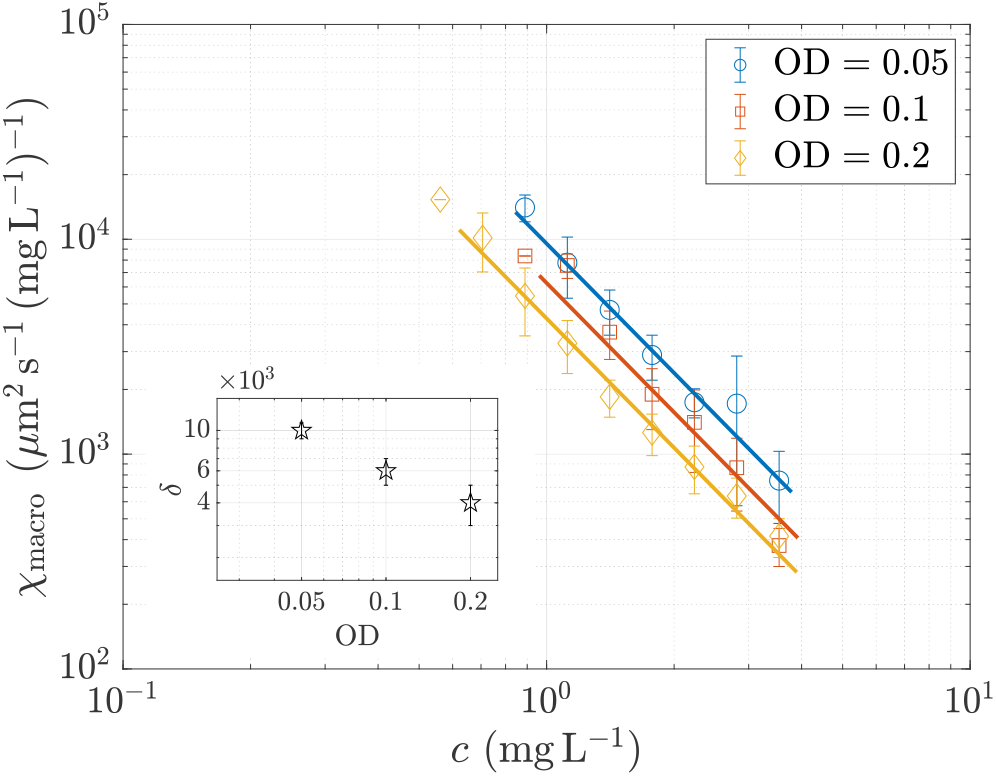
Aerotactic response *χ*_macro_ ≃ *v*_*x*_*/*(*∂c/∂x*) where *∂b/∂x* 0, plotted as a function of the oxygen concentration *c* for different initial bacterial concentrations: OD = 0.05 (blue circles), OD = 0.1 (orange squares) and OD = 0.2 (yellow diamonds). The lines show the best fit *χ*_macro_ = *δ/c*^2^ for each OD, the coefficient *δ* being plotted as a function of the OD in the inset. For *c >* 4 mg L^*−*1^, the aerotactic velocity is too close to zero for a reliable measurement of *χ*_macro_. result from a secondary negative chemotactic response lowering the effect of the primary positive aerotactic response.

## IV. AEROTACTIC RUN TIME MODULATION

Measuring the aerotactic coefficient *χ*(*c*) from the mean drift velocity *v*_*x*_ is inherently limited by the effect of diffusion, which restricts our measurements to regions where the diffusive flux vanishes, for *∂b/∂x* ≃ 0. To circumvent this limitation, we now investigate the aerotaxis at the microscopic level, with the aim to characterise the modification of the swimming pattern statistics due the oxygen gradient.

For each track, we define a run as a continuous segment between two tumbles [Fig. 2(g)], such that the swimming direction changes abruptly, | *δθ/δt* | *> ω*_*c*_, with *ω*_*c*_ ≃ 4 rad s^*−*1^ as threshold. Runs defined in this way consist of nearly straight lines with a large coherence length. For each run *j*, we define the mean velocity **V**_run,*j*_, the mean angle *θ*_run,*j*_ with respect to the *x* axis and the run time *τ*_run,*j*_.

To characterise the aerotactic bias of the run times, we partition the runs in two sets specified by the sign of cos *θ*_run,*j*_, and define the corresponding run times as 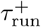 (bacteria swimming towards the oxygen source) and 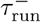 (bacteria swimming in the opposite direction). Figure 7 shows the distribution of these run times, with and without aerotaxis. At early time [Fig. 7(a)], without significant oxygen gradient, 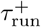 and 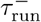 follow the same Poisson distribution, 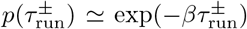, with *β* ≃ 1.47 s^*−*1^. At larger time [Fig. 7(b)], in the migration band, the two distributions still decay exponentially but with different decay rates, confirming that bacteria swim longer when moving towards the oxygen source.

**Figure 7.**
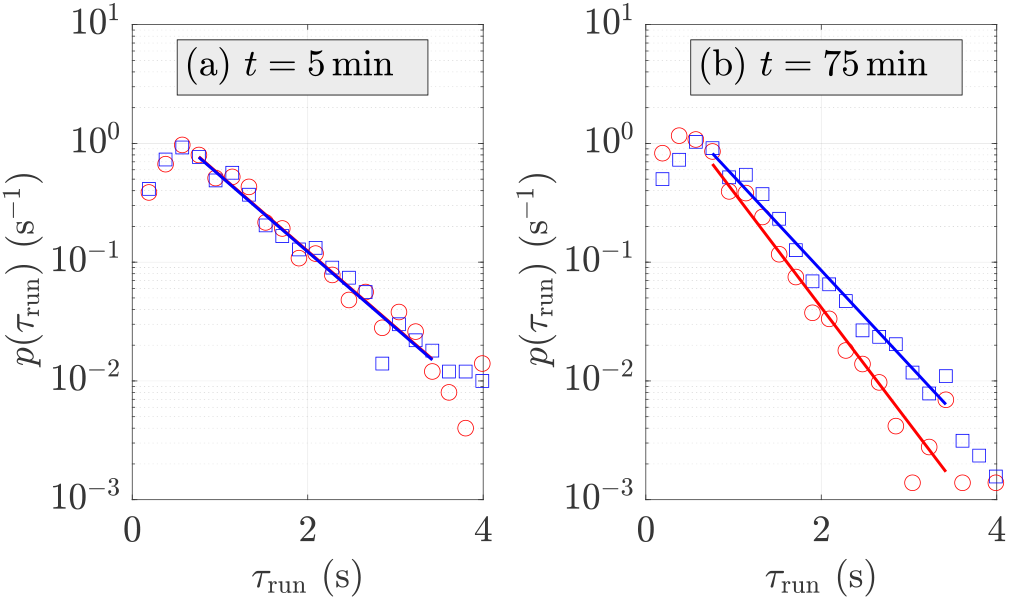
Distributions of run times 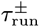 measured at OD = 0.05, *x* ≃ 4 *−* 5 mm, at times *t* = 5 min (a), before the aerotactic front, and *t* = 75 min (b), in the aerotactic front. Blue: 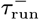, conditioned on cos *θ*_run,*j*_ *<* 0. Red: 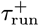, conditioned on cos *θ*_run,*j*_ *>* 0. The lines show exponential fits 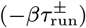, with (a) *β*^*±*^ ≃ 1.47 s^*−*1^, and (b) *β*^*−*^ ≃ 2.24 s^*−*1^, *β*^+^ ≃ 1.83 s^*−*1^.

To further quantify this effect, we compute, for each time *t* and location *x* along the capillary, the mean run duration conditioned on the mean run angle,

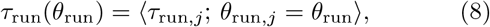

with ⟨ · ⟩ the average over the runs *j*. We plot this mean run time *τ*_run_ as a function of cos *θ*_run_ in Fig. 8, both without and with oxygen gradient. We observe a welldefined linear trend,

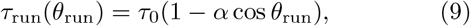

with *α* the run time modulation coefficient, and *τ*_0_ the mean run time at cos *θ*_run_ = 0. Such a linear dependency of *τ*_run_ with cos *θ*_run_ comes from the fact that bacteria are sensitive to the time variation of the local oxygen gradient sampled along their trajectory, *dc/dt* ≃ **V** · **∇***c* = *V* cos *θ ∂c/∂x* (see Appendix D). Without oxygen gradient, we have *α* ≃ 0 (isotropic swimming), whereas in the oxygen gradient, runs towards the oxygen source are (1 + *α*) longer than *τ*_0_ while runs away from the oxygen source are (1 − *α*) shorter than *τ*_0_.

**Figure 8.**
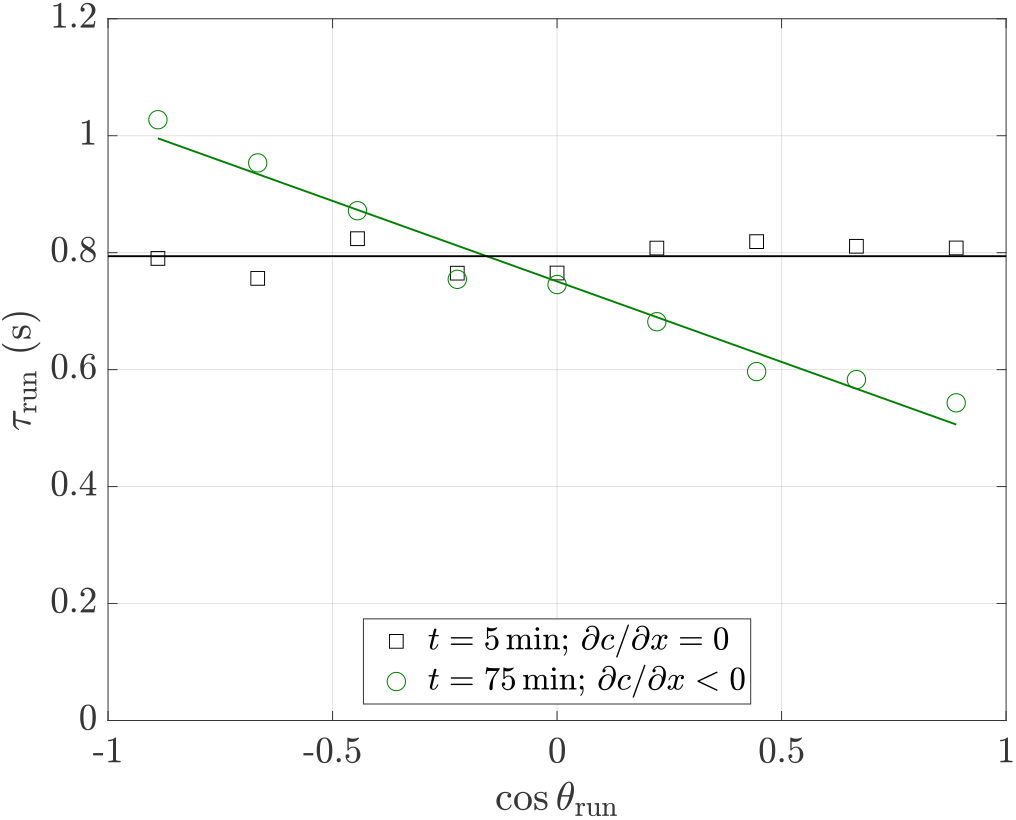
Average run time *τ*_run_ as a function of cos *θ*_run_ (same data as in Fig. 7). In the aerotactic band, the run time increases for bacteria swimming towards the oxygen source (cos *θ*_run_ = *−*1). The green line shows the linear fit using Eq. (9) with *α* 0.28, and the black line shows the case *α* = 0 (no aerotaxis).

From the run statistics, we extract the run time modulation coefficient *α* at each time, for each location *x* along the capillary chamber, and for the three bacterial concentrations (OD = 0.05, 0.1 and 0.2). The time evolution of *α*(*x*), plotted in Fig. 9 for OD = 0.2, shows a clear drift of the aerotactic response towards the oxygen source as the migration band proceeds, which is consistent with the measured drift velocity − *v*_*x*_(*x*) in Fig. 4. Assuming that the swimming velocity *V*_0_ and the mean run time *τ*_0_ do not depend on *θ*_run_, using Eq. (9) and averaging over all swimming directions yield (see Appendix D)

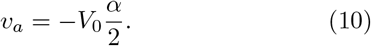

**Figure 9.**
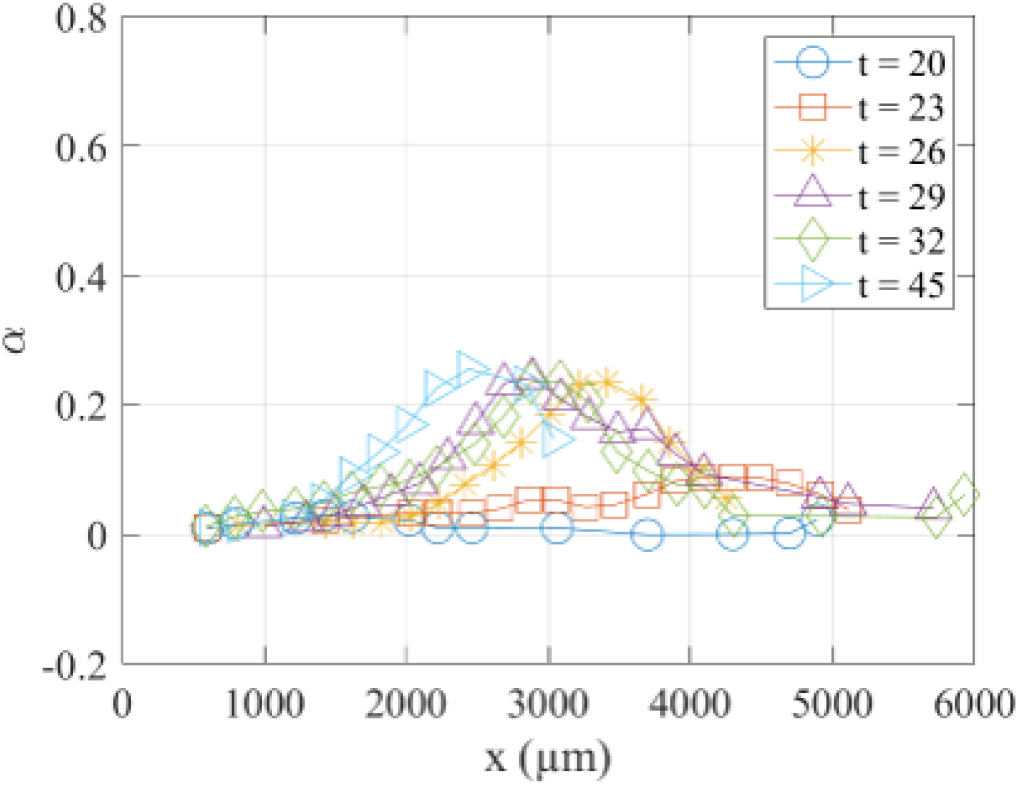
Run time modulation coefficient *α* as function of the distance from the oxygen source at different times during the migration of the bacterial band. Same data as in Fig. 4.

Interestingly, this estimate of *v*_*a*_ from *α* is expected to hold everywhere, not only at the maximum of the migration band where the diffusion contribution vanishes. This is clearly visible by comparing Figs. 4 and 9: while *v*_*x*_ drops to 0 at long time because aerotaxis is balanced by diffusion in the asymptotic state, a strong aerotactic response is still present at the bacterial scale in the run time modulation coefficient *α*.

We finally plot in Fig. 10 the aerotactic coefficient [Eq. (1)] computed using Eq. (10), which we call *χ*_micro_. This coefficient is again compatible with the scaling *c*^*−*2^ for all bacterial concentrations. Since we are not limited here to *∂b/∂x* ≃ 0, the range of oxygen concentration where *χ*_micro_ can be computed is extended to lower oxygen concentrations, down to *c* ≃ 0.4 mg L^*−*1^, i.e., over nearly one decade in *c*.

**Figure 10.**
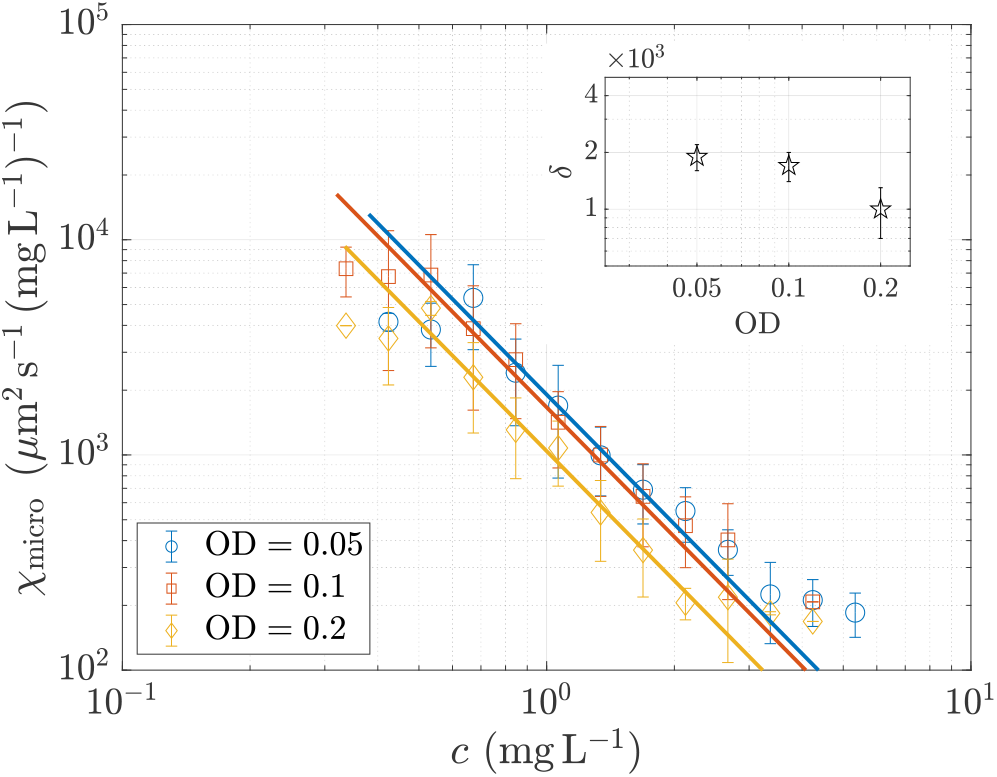
Aerotactic response *χ*_micro_ ≃ *− αv/*(2*∂c/∂x*) plotted as a function of the oxygen concentration *c* for different bacterial concentrations: OD = 0.05 (blue circles), OD = 0.1 (orange squares) and OD = 0.2 (yellow diamonds). The lines show the best fit *χ*_micro_ = *δ/c*^2^ for each OD, the coefficient *δ* being plotted as a function of the OD in the inset.

Although the aerotactic responses computed at the population scale (Fig. 6) and at the bacterial scale (Fig. 10) both display the same scaling *c*^*−*2^, a systematic shift is observed between these two estimates, with *χ*_micro_ ≃ 0.3 *χ*_macro_. This discrepancy may be attributed to several biases in the segmentation of tracks into runs. First, since only the two-dimensional projection of tracks can be measured, runs nearly normal to the measurement plane are perceived as low-velocity erratic motions and can be interpreted as tumbles. Second, the measurements are limited both in depth (the depth of field is 30 μm) and acquisition time (image sequences are limited to 5 s to allow for a fast scan along the *x*-direction). Longer tracks are therefore truncated, resulting in a large number of small tracks without tumbles. This leads to an over-representation of short runs, and hence an underestimation of *α*. The aerotactic response *χ*_micro_ computed from the run time modulation therefore underestimates the true aerotactic response, but its overall scaling *c*^*−*2^ remains unaffected by this bias.

## V. CONCLUSION

In this paper, we use an original setup allowing for simultaneous measurements of oxygen concentration and bacteria tracking to investigate the scaling of the aerotactic response. Besides giving us access to multiple oxygen conditions, this setup allows us to quantitatively monitor bacteria migrating to an oxygen source, as in natural environments. The aerotactic response *χ* computed both macroscopically, from the concentration profiles of bacteria, and microscopically, from the bias in run times, yields consistent results in the form *χ* ∝ *c*^*−*2^. To our knowledge, this is the first direct measurement of *χ*(*c*).

The scaling *χ* ∝ *c*^*−*2^ is consistent with the models based on the biochemistry of bacterial membrane receptors [31, 34, 35] given by Eq. (4) for a dissociation constant *K*_*d*_ much smaller than the typical oxygen concentration. It differs from the “log-sensing” scaling *χ* ∝ *c*^*−*1^ reported in recent experiments using stationary oxygen gradients [4, 37] with other bacteria. This variability may originate from a specific aerotactic response of the different bacterial species, which exhibit different oxygen receptors or swimming patterns (“run-and-tumble”, “run-reverse” or “run-reverse and flick” [47, 49]).

The simultaneous mapping of oxygen concentration using a fluorescent O_2_-sensor together with the bacteria tracking is a very promising approach. It would be of prime interest to extend this study to more complex chemical stimuli, when aerotaxis competes with other chemotaxis, or to the transport of bacterial populations through complex media like natural soils, as found in bacterial ecosystems.

## Appendix A Oxygen measurements

Oxygen concentration maps are obtained from fluorescence microscopy images of Ruthenium, an organometallic complex quenched by oxygen. Ruthenium complex [ruthenium-tris(4,7 diphenyl-1, 10-phenanthroline) dichloride, also named Ru(dpp)_3_Cl_2_ or Ru(dpp)] is encapsulated in phospholipidic micelles (referred to as Ru-micelles), in order to make it soluble in water and bio-compatible [6, 41]. We did not observe any influence of Ru-micelles on the bacterial motility.

Fluorescence intensity is averaged over the *y*-direction and then converted into O_2_ concentrations using the Stern-Volmer relation

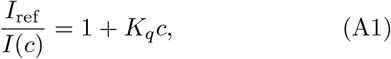

where *I*(*c*) is the fluorescent intensity at oxygen concentration *c, I*_ref_ is the intensity without oxygen, and *K*_*q*_ is the quenching constant. *I*_ref_ is taken as the highest intensity for each experiment. This reference level was confirmed to be equal to *c* = 0 mg L^*−*1^ when measured with an electrochemical O_2_-probe in a closed sample where bacteria consumed the available oxygen. Figure 11 shows the temporal evolution of the oxygen concentration in two bacterial suspensions of different concentrations enclosed in a glass capillary with no oxygen supply.

**Figure 11.**
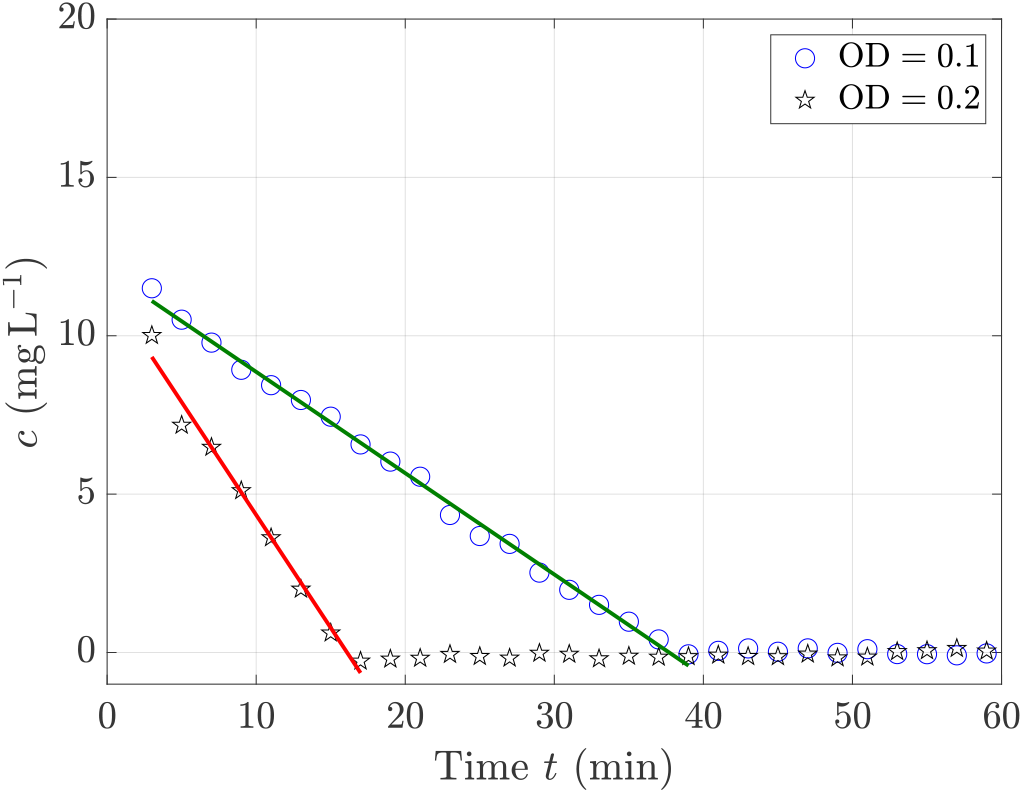
Temporal evolution of the dioxygen concentration *c* in a bacterial suspension of OD = 0.1 (blue circles) and OD = 0.2 (black stars). The oxygen initially dissolved is consumed at a steady rate by *B. contaminans* bacteria. The concentration is deduced from the fluorescence intensity thanks to the Stern-Volmner relation [Eq. (A1)]. The black line shows the linear fit *c*(*t*) = *c*_0_ *− k*_*c*_*t* with *c*_0_ = (11.8 *±* 3.0) mg L^*−*1^ and a consumption rate *k*_*c*_ of 0.32 and 0.71 mg L^*−*1^ min^*−*1^.

*K*_*q*_ was determined by measuring the fluorescent intensity of a solution containing no oxygen (a solution saturated of *B. contaminans* bacteria), and comparing it to the intensity level of a solution with a known oxygen concentration. We find *K*_*q*_ = 0.21 L mg^*−*1^, which is of the same order of magnitude of values reported for similar Ruthenium-based dyes [4, 5, 50–53].

## Appendix B “Run-reverse” swimming of *B. contaminans*

*B. contaminans* is a motile bacterium exhibiting a large dispersion of velocities. Its velocity (projected on the microscope plane) ranges from 0 up to 60 μm s^*−*1^, with a mean value around 20-25 μm s^*−*1^. A particular feature of *B. contaminans* is its “run-reverse” swimming (illustrated in Fig. 2), similar to some marine bacteria [46, 47].

To quantify this run-reverse motion, we measure the turning angle *δθ* between two successive runs, defined as *δθ* = *θ*_run,*j*+1_ − *θ*_run,*j*_ between runs *j* and *j* + 1. Its distribution, plotted in Fig. 12, shows that a majority of tumbles correspond to reverses. We obtain a mean value of cos *δθ* ≃ −0.34, i.e., turning angles much sharper than that of *E. coli* (⟨cos *δθ*⟩≃ 0.33 [45]).

**Figure 12.**
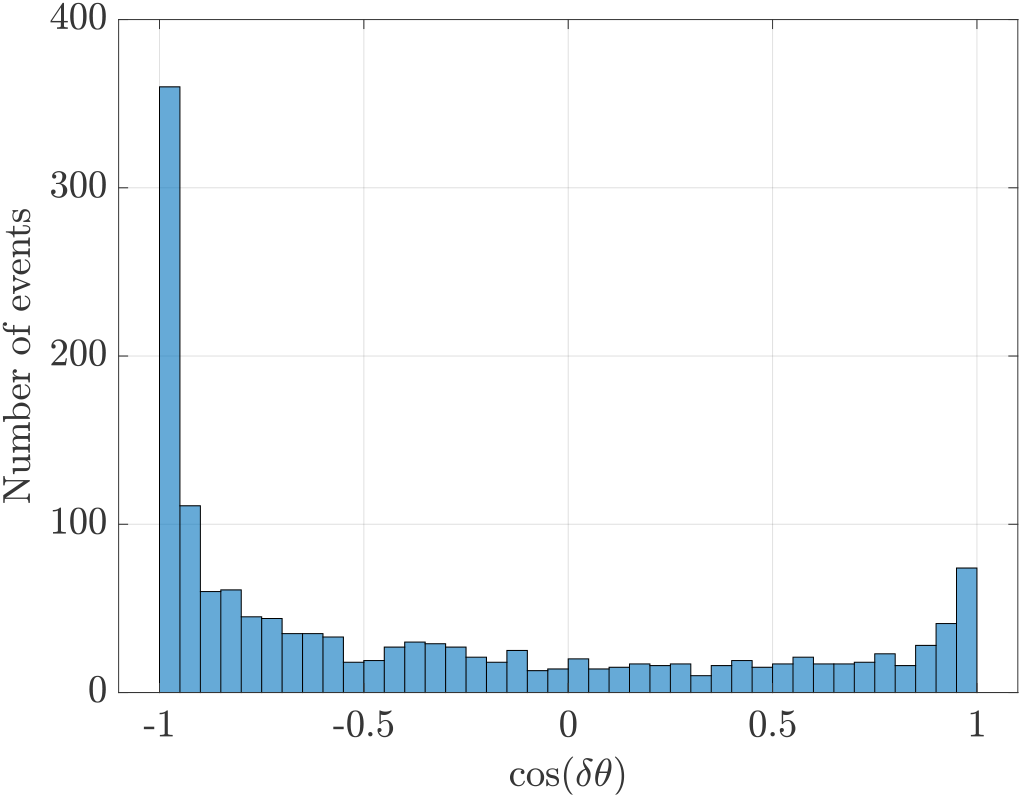
Distribution of the turning angle *δθ* between two successive runs. The distribution is peaked at cos *δθ* = *−*1, corresponding to reverses.

## Appendix C Diffusion coefficient

To measure the diffusion coefficient *μ*, we compute the velocity correlation function [47, 54, 55]

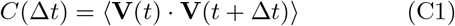

averaged over all the bacterial tracks. An example of *C*(Δ*t*) is shown in Fig. 13. We fit the exponential decrease as

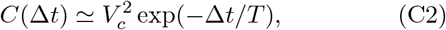

with 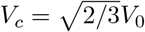, and *V*_0_ the mean swimming velocity, from which we compute the diffusion coefficient as 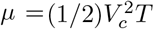. An average over numerous experiments yields *μ* ≃ (45 0 ± 100) μm^2^ s^*−*1^.

**Figure 13.**
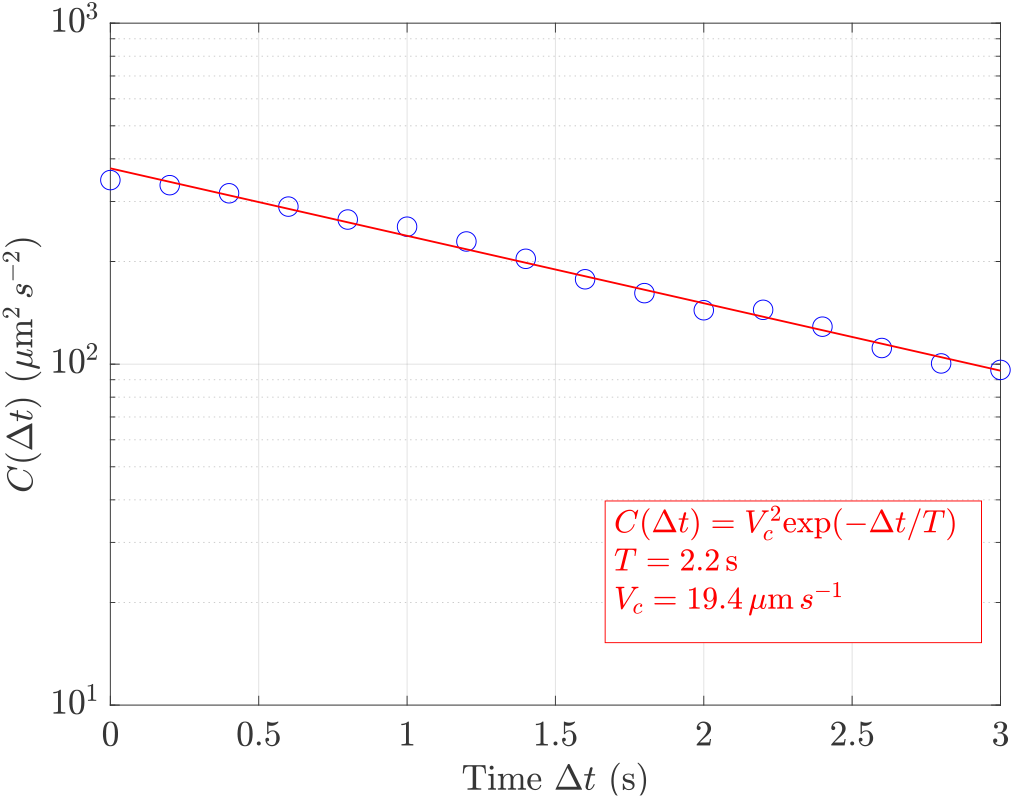
Velocity correlation function *C*(Δ*t*) = ⟨**V**(*t*) *·* **V**(*t* + Δ*t*) ⟩ for a *B. contaminans* suspension of OD = 0.05. The red line shows the exponential fit [Eq. (C2)], with *V*_*c*_ = 19.4 μm s^*−*1^ and *T* = 2.2 s.

## Appendix D Aerotactic run time modulation

In this appendix, we examine the aerotactic bias in the run time distribution. We first consider nonaerotaxict run-and-tumble (homogeneous oxygen concentration). The probability *p*(*τ*_run_) that a bacteria makes a run of duration *τ*_run_ is

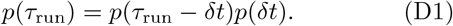

Noting *β* the probability per unit time of a tumble, we have *p*(*δt*) = 1 − *βδt*, and Eq. (D1) becomes

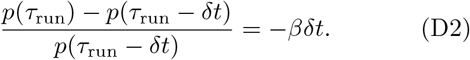

The probability *p*(*τ*_run_) is then given by

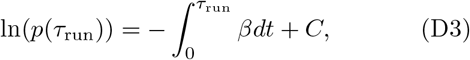

where *C* is an integration constant defined such that 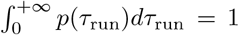. For constant *β*, the solution of Eq. (D3) is a Poisson law, *p*(*τ*_run_) = *β* exp(−*βτ*_run_), and the average duration of a run is

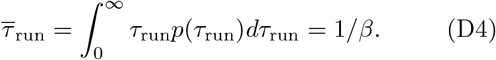

We consider now the run time distribution in the presence of an oxygen gradient. We derive the average of the runs of bacteria moving up 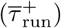 or down 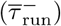 the oxygen gradient under the assumptions that (i) the frequency *β* depends only on the local oxygen concentration *c*, and (ii) the bacteria swim at a constant velocity *V*_0_.

The probability to make a run of duration *τ*_run_ is still given by Eq. (D3) but with a frequency *β* = *β*[*c*(*t*)] that depends on the local oxygen concentration. The average duration of the runs of the bacteria moving up or down the gradient are

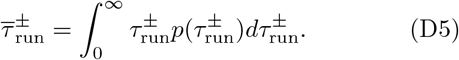

A Taylor expansion of *β*[*c*(*t*)] around its initial value at *t* = 0 gives

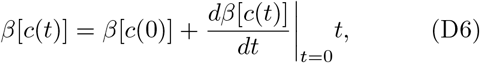

Yielding

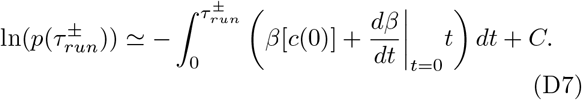

For a constant swimming velocity, the time derivative of *β* writes

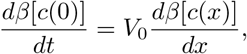

where *x* is the bacteria position at time *t* = 0. We define the run-time modulation coefficient as

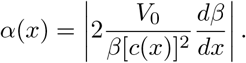

In the limit of small *α*, Eq.(D5) reduces to

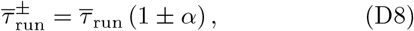

where 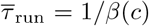 is the average run duration of bacteria moving in an homogeneous oxygen concentration *c* [Eq. (D4)]. We note that Eq. (D8) can be written in a similar way than proposed by Rivero *et al*. [31] and Berg and Brown [44],

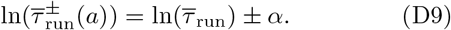

The calculation can be generalised by considering runs not aligned with the gradient, but making an angle *θ*_run_ with respect to the gradient,

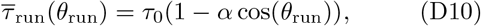

where *τ*_0_ is the average run time measured normal to the gradient (for cos *θ*_run_ = 0). Here, we have *θ*_run_ = *π* for runs moving up the gradient.

The aerotactic velocity can be written

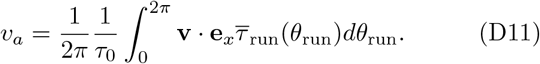

Assuming *τ*_0_ and *V*_0_ do not depend on the run angle, we obtain

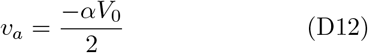

by combining Eqs. (D10) and (D11).

## Appendix E Concentration profiles

**Figure 14.**
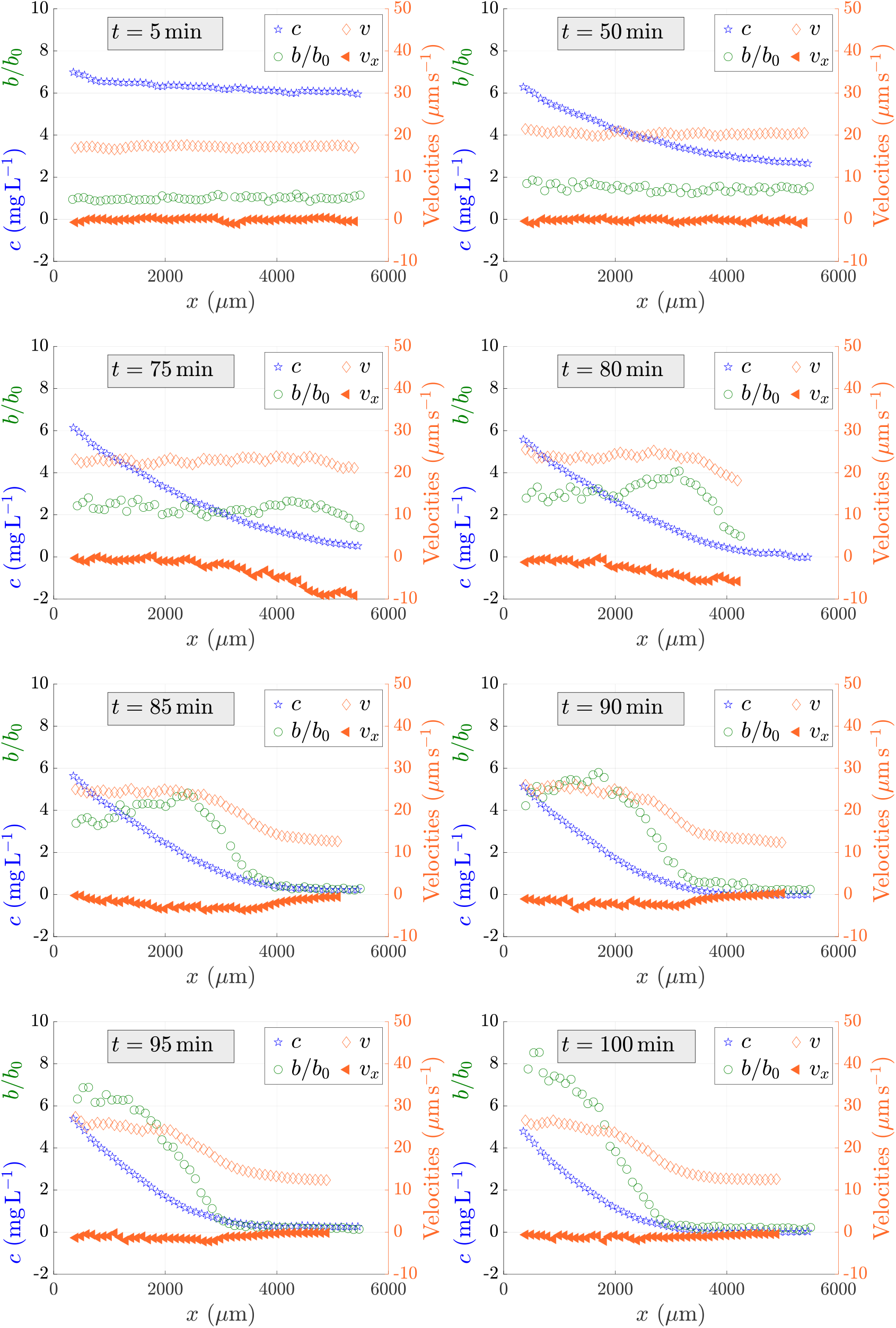
Variations of the local density of bacteria *b*(*x*) (green circles), oxygen concentration *c*(*x*) (blue stars), magnitude of the swimming velocity of bacteria *v*(*x*) (empty orange diamonds) and drift velocity along the gradient *v*_*x*_(*x*) (filled orange diamonds) as functions of the distance from the oxygen source at four different times during the migration of the bacterial band. The capillary is initially filled with an homogeneous suspension of bacteria at a concentration OD = 0.05. Time is counted from the sealing of the capillary.

**Figure 15.**
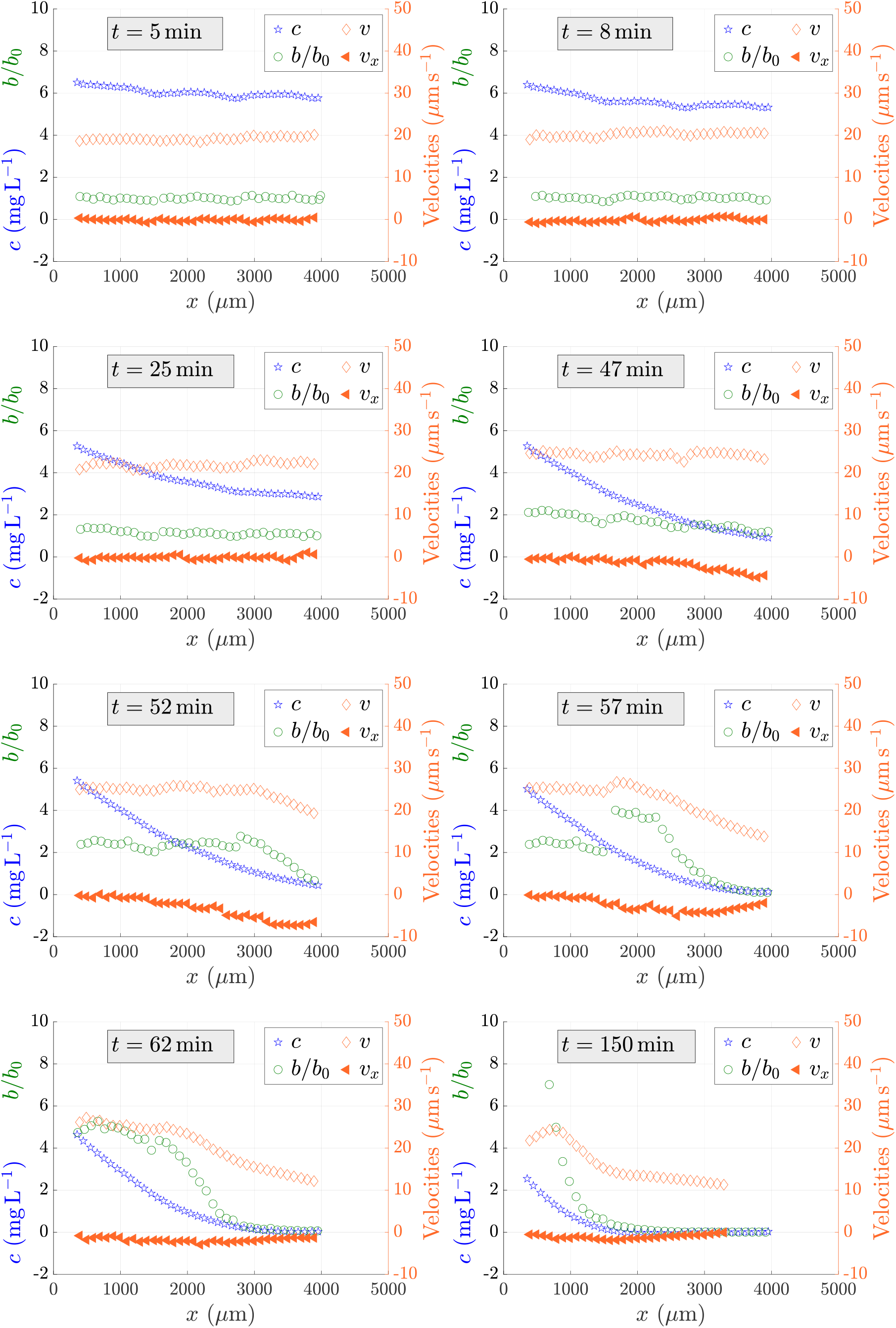
Same as in Fig. 14 for OD = 0.1.

**Figure 16.**
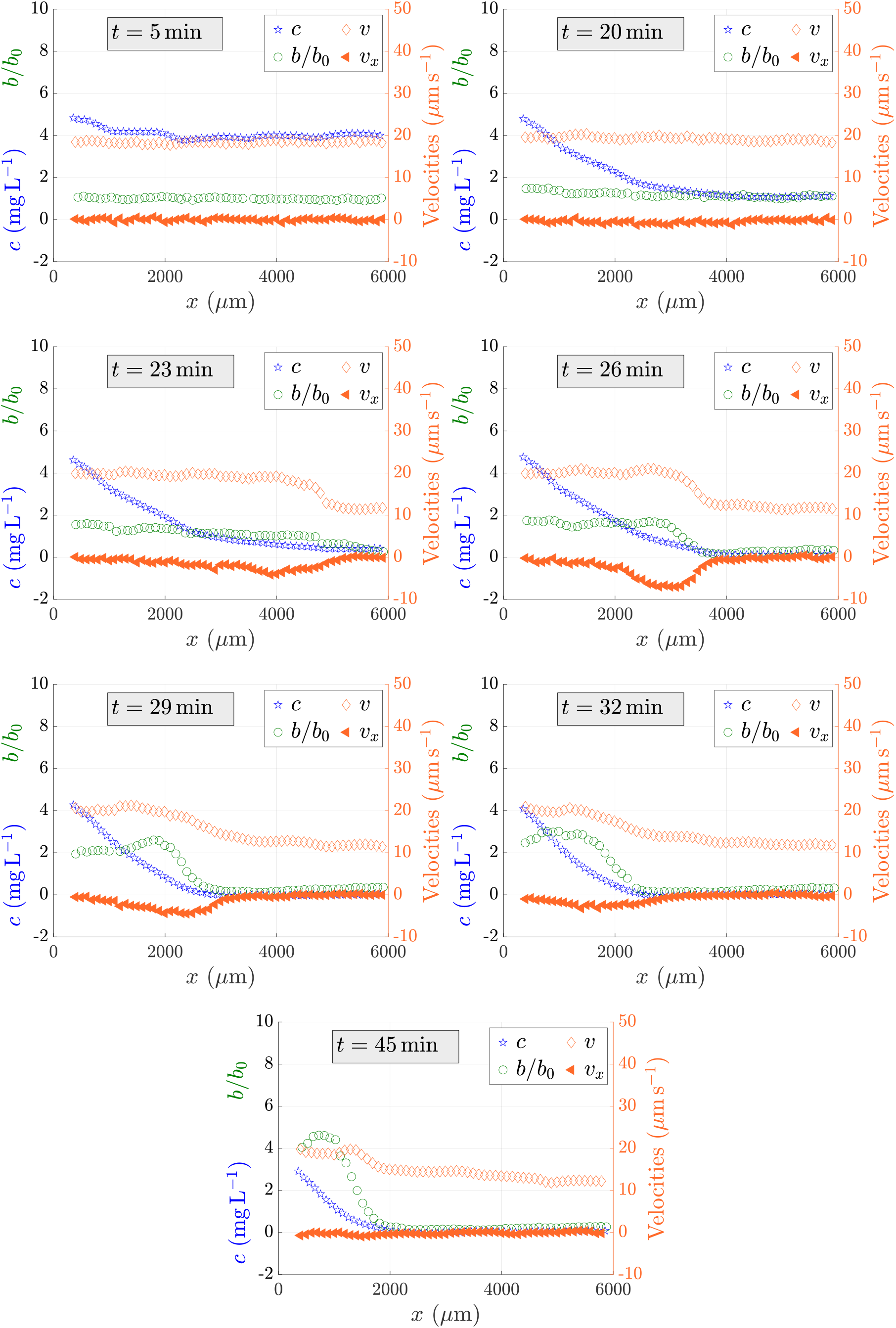
Same as in Fig. 14 for OD = 0.2.

